# SpaRC: Scalable Sequence Clustering using Apache Spark

**DOI:** 10.1101/246496

**Authors:** Lizhen Shi, Xiandong Meng, Elizabeth Tseng, Michael Mascagni, Zhong Wang

## Abstract

Whole genome shotgun based next generation transcriptomics and metagenomics studies often generate 100 to 1000 gigabytes (GB) sequence data derived from tens of thousands of different genes or microbial species. *De novo* assembling these data requires an ideal solution that both scales with data size and optimizes for individual gene or genomes. Here we developed a Apache Spark-based scalable sequence clustering application, <u>Spa</u>rk<u>R</u>ead<u>C</u>lust (SpaRC), that partitions the reads based on their molecule of origin to enable downstream assembly optimization. SpaRC produces high clustering performance on transcriptomics and metagenomics test datasets from both short read and long read sequencing technologies. It achieved a near linear scalability with respect to input data size and number of compute nodes. SpaRC can run on different cloud computing environments without modifications while delivering similar performance. In summary, our results suggest SpaRC provides a scalable solution for clustering billions of reads from the next-generation sequencing experiments, and Apache Spark represents a cost-effective solution with rapid development/deployment cycles for similar large scale sequence data analysis problems. The software is available under the Apache 2.0 license at https://bitbucket.org/LizhenShi/sparc.

## Introduction

Whole genome shotgun sequencing (WGS) using next-generation sequencing technologies followed by *de novo* assembly is a powerful tool for *de novo* sequencing large eukaryote transcriptomes (reviewed in [22]) and complex microbial community metagenomes (reviewed in [37]) without reference genomes. Because of the stochastic sampling nature associated with WGS and the presence of sequencing errors, it is necessary for the reads to cover a single gene or genome many times (coverage), typically 30x-50x, to ensure high quality *de novo* assembly [2]. Unlike in single genome sequencing projects where the majority of the genomic regions are equally represented, in transcriptome and metagenome sequencing projects, different species of transcripts or genomes may have very unequal representation, up to several orders of magnitude in abundance difference[16, 23]. To obtain a good assembly covering low abundant species one would sequence the population at much higher depth than single genome projects. As in practice it is difficult to precisely estimate the required sequencing depth without knowing the community structure, sequencing large transcriptomes and complex metagenomes often generates as much data as the budget allows, producing 100-1000 GB of sequence data or more [15] [33]. The largest project so far is the Tara Ocean Metagenomics project where 7.2 Tera bases (Tb) was generated [35].

Since current NGS technologies are not able to read the entire sequence of a genome at once, genomes are broken into small DNA/RNA fragments followed by massive parallel high-throughput sequencing. Different technologies produce sequence reads that vary in length. For example, Illumina technology [17] typically generates about 150 base pair(bp) per read, comparing to 100 bp to 15,000 bp by Pacific Biosciences [28]. Assembling these reads to obtain genomes is challenging due to it is both a compute and memory-intensive problem, and this challenge grows exponentially with the size of dataset. Further, the *de novo* assembly problem is compounded by the quality of sequencing data and the presence of certain genetic complexity (repeat elements, species homology, etc). For a comprehensive review of *de novo* assembly algorithms please refer to [25]. Assembling a large dataset as a whole also requires efficient computing, and these assemblers use either multiple processes on a shared memory architecture (MetaSPAdes[26], MEGAHIT[19], etc) or MPI to distributed on a cluster[11]. The shared memory approach is very hard to scale up to exponentially increased NGS data size. In addition, these assemblers try to tackle the problem as a whole and is not able to produce optimized results as different transcripts or genomes may need individualized optimal parameter settings.

Our work was initially inspired by a “divide-and-conquer” approach presented by DIME [13]. DIME first clusters reads based on their overlap, then assembles them separately. It was implemented using Apache Hadoop [4] platform and in theory should scale to large data sets. In practice, however, Hadoop-based implementation has very poor computing efficiency, making it expensive to run on commercial cloud providers. Further, Hadoop-based solutions often produce much larger intermediate files than the input files during the assembly, making them harder to scale to very large datasets.

Recently Apache Spark [5] has overtaken Hadoop in the big data ecosystem due to its fast in-memory computation. Spark has been successfully applied to several genomics problems such as [39, 18, 7, 24, 8]. In this paper we developed a new algorithm based on Spark’s GraphX library, called Spark Read Clustering (SpaRC), for parallel construction and subsequent partition of NGS sequence reads. For scalability and computing efficiency SpaRC implemented several heuristics: 1) the pairwise similarity between sequences (edge weight) is estimated by the number of shared k-mers; 2)To control the explosion of edges as data complexity grows, SpaRC sets a max number of edges a vertex can have for each shared k-mer; 3) it adopts an overlapping community detection algorithm - Label Propagation Algorithm (LPA) [30] - to partition the read graph, therefore avoiding the formation of very large partitions due to repetitive or other shared genetic elements between species. We report clustering accuracy and computing performance on both transcriptomic and metagenomic datasets from both short read (Illumina) and long read (PacBio) sequencing platforms.

## Methods

### Algorithm Overview

**SpaRC** is a generic sequence read clustering algorithm as it is designed for clustering both short and long reads. It first computes the number of shared k-mers between a pair of reads to approximate their overlap, and then builds an undirected read graph followed by graph partitioning to achieve read clustering. Specifically, it contains four modules: K-mer Mapping Reads, Graph Construction and Edge Reduction, Graph Partition, and Sequence Retrieval. We describe each of these modules in details as the following.

### K-mer Mapping Reads (KMR)

Given a set of sequence reads, KMR splits them into k-mers according to a predefined k-mer length and only keeps distinct k-mers for each read to avoid low-complexity sequences. KMR keeps track of each k-mer and the reads containing it. The length of k-mer (*k*) is a parameter to control the sensitivity and specificity of read overlap detection. Shorter k-mers result in more sensitivity but less specificity and vice versa. The ideal k-mer size may depend on sequencing platform, read depth, and sequence complexity.

Generally k-mers appear in only a single read are derived from either sequencing errors or rare molecules, and they take up a large fraction of the total k-mers but are not useful for computing read overlap, therefore they are filtered out. KMR allows users to specify customized filtering criteria (*min_kmer_count* and *max_kmer_count*) for more stringency.

### Graph Construction and Edge Reduction (GCER)

GCER constructs a read graph where a node is a read and an edge links two nodes if they share k-mers. Some nodes, if derived from repetitive elements, homologous genes among species or contamination, can have extremely high number of edges (degrees). GCER sets the maximum degree of any vertex for each shared k-mer in a graph (*max degree*) as a parameter to reduce unnecessary computation.

After all the vertices and edges are generated, GCER then merges the edges having the same source and destination and filters out those edges with the number of shared k-mers less than the specified parameter *min shared kmers.*

### Graph Partition

SpaRC provides two algorithms for iterative graph partition, Label Propagation Algorithm (LPA) [30] (by default) and Connected Components (CC) [9]. As repetitive elements and homologous genetic elements shared between different molecules/genomes create “overlap communities”, in practice LPA in general works much better than CC because LPA allows the resolution of overlap communities. For dataset with very low sequencing coverage CC may be useful.

### Sequence Retrieval (AddSeq)

In the above modules reads are represented by numeric IDs to save memory and storage. Once the clusters are formed, AddSeq retrieves the sequences and get them ready for downstream parallel assembly with a choice of an assembler.

#### Algorithms

~~~
For each read r in the read s e t R:
    Generate distinct kmer–read tuples

Group the tuples by k–mer and generate kmer–reads pairs (KR)
Filter KR by only keeping pairs over lapping between *min_kmer_count*
   and *max_kmer_count*

For each list of reads in KR:
    Generate pairwise edges (reads as nodes)

For each node in the edges:
    If the node degree > *max_degree*, sample *max_degree* edges

Count distinct edges and generate edge–count pairs (EC)
Filter EC to only keep pairs whose count is more than *min_shared_kmers*
Generate graph *g*0 with the edges in EC

If clustering algorithm ɖ is CC:
    Generate the connected components of *g*0.
    For each connected component, add the connected
     component to the set of read clusters Ω.
else if ɖ is LPA:
    Run label propagation step for m iterations
    Group the nodes (reads) by its labels
    For each reads group, add the group to Ω
~~~

### Hardware and Software Environment

We implement the above algorithm using the Scala (Scala 2.11.8) functional programming language [36]. Performance was tested on two closely matched cloud environments, 20 nodes Open Telekom Cloud (OTC) and Amazon’s Elastic MapReduce (EMR, emr-5.9.0). For these two clusters, one node is used as the master and all other nodes are used as workers. Configuration details are shown in Table 1.

**Table 1:**
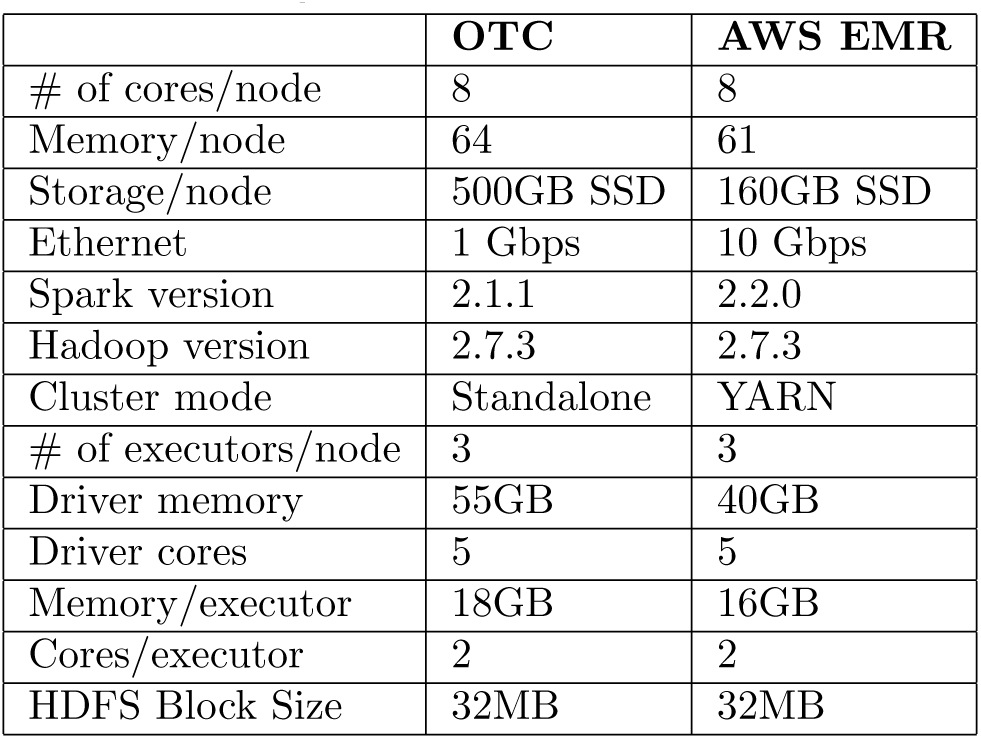
Configuration for OTC and AWS EMR.

|  | OTC | AWS EMR |
| --- | --- | --- |
| # of cores/node | 8 | 8 |
| Memory/node | 64 | 61 |
| Storage/node | 500GB SSD | 160GB SSD |
| Ethernet | 1 Gbps | 10 Gbps |
| Spark version | 2.1.1 | 2.2.0 |
| Hadoop version | 2.7.3 | 2.7.3 |
| Cluster mode | Standalone | YARN |
| # of executors/node | 3 | 3 |
| Driver memory | 55GB | 40GB |
| Driver cores | 5 | 5 |
| Memory/executor | 18GB | 16GB |
| Cores/executor | 2 | 2 |
| HDFS Block Size | 32MB | 32MB |

### Datasets

We tested the performance of SpaRC on both simulated and real world datasets. A maize sequence dataset we generated previously from [23], and the cow rumen metagenome dataset [14], from which we generated subsets of 1 to 100 (GB) in fastq, for testing scalability. A mock dataset containing 26 genomes is used to verify accuracy. Three long read transcriptome datasets were provided by PacBio. The datasets are described in Table 2.

**Table 2:** Metrics of test datasets.

| Dataset | # Species | Read Length (bp) | Size(GB) |
| --- | --- | --- | --- |
| PacBio1 (Human Alzheimer brain transcriptome) | High | 300 - 30816 | 3.8 |
| PacBio2 (Human UHRR + synthetic RNA, 2Cell) | High | 62 - 14621 | 1 |
| PacBio3 (Human UHRR + synthetic RNA, 3Cell) | High | 54 - 14833 | 1.8 |
| Maize transcriptome | High | 151X2 | 4 |
| Mock metagenome | Low (26) | (90-150)x2 | 15 |
| Cow Rumens metagenome | Medium | 100x2 | 100 |

## Results

### SpaRC clustering accuracy

In order to measure the clustering performance of SpaRC, we used two sets of real world data sets with “known answers” and ran SpaRC to obtain clusters.

The first dataset is derived from human Alzheimer whole brain transcriptome sequenced by PacBio consisting of 1,107,889 full-length transcript sequences. The transcript sequences were first clustered together based on an isoform-level clustering algorithm [12], then the consensus sequence from each cluster were mapped back to the human genome to identify which loci it came from. Reads coming from clusters where the mapped genomic location overlap by at least 1 bp are considered to be from the same loci. This is the theoretical limit for overlap-based clustering algorithms. The second data set is two million Illumina short metagenome reads (150bp) sampled from a mock microbial community consisting of 26 genomes described previously [34]. Clusters are defined similarly as above for the PacBio transcriptome data set.

By comparing the SpaRC clusters to “known answers” in the above two datasets, we measured SpaRC’s performance by cluster purity, and cluster completeness. Here cluster purity is defined as the percentage of reads belonging to the dominant known cluster for each SpaRC cluster, and completeness is defined as the maximum percentage of reads from a known cluster could be captured by a SpaRC cluster. It is worth noting that cluster completeness will be an underestimation of the true cluster completeness as the “known answers” are overestimation as described above. As the sensitivity of overlap detection is heavily influenced by the read length, in the Illumina metagenome dataset we joined the reads in a pair that are pair-end sequenced to double the read length for clustering.

In both experiments SpaRC clustered the majority of the reads (PacBio: 82.65%, Ilumina: 98.3%), and generated very pure clusters ( Fig 1 A,E). For the impure read clusters in the both datasets, the contamination events seem to be relative low, as their purity increases with cluster size (Fig 1 B,F).

**Figure 1.**
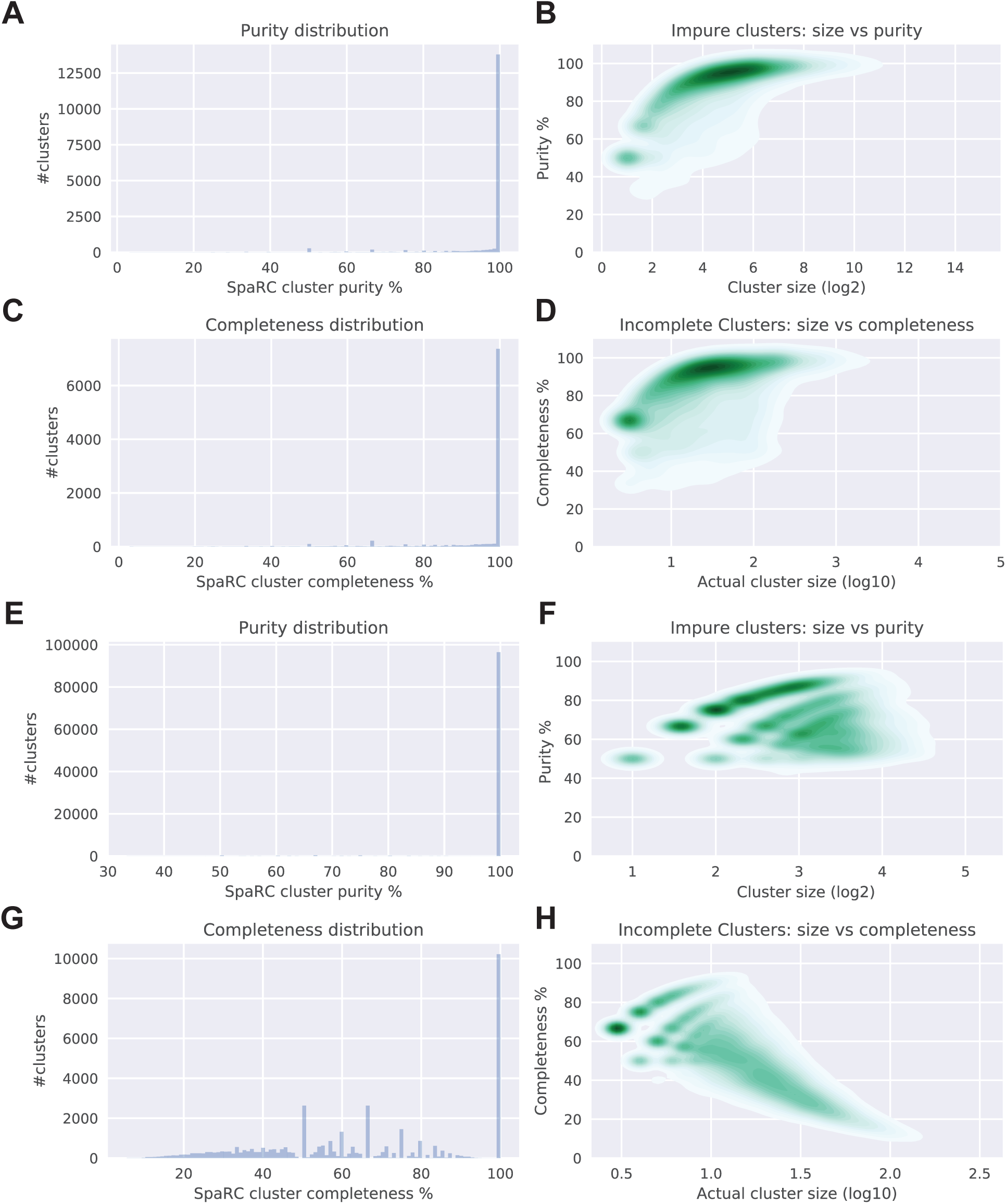
SpaRC’s clustering performance on long and short reads. A-D) Pacbio transcriptome and E-H) Illumina metagenome.

Clustering long reads achieved a much higher completeness than short reads, with many more clusters that have completeness ≥90% (84.88%, n=9,578, Fig 1 C), comparing to short read clusters (37.19%, n=37,879), Fig 1 G). For the long read transcriptome dataset, the completeness improves as cluster size is getting bigger, suggesting more copies of a transcript increases the chance of finding overlap. For the short read metagenome dataset, larger clusters tend to have lower completeness. As the copy number of each genome is a constant and larger clusters translate to larger genome regions, they are more prone to be broken into smaller clusters due to loss of some overlaps.

We also tested whether or not completeness would get worse if the read pairs in the short read dataset were not joined. This indeed is the case, as clusters that have completeness ≥90% is decreased to 6.08% and more small clusters are produced (n=42,181).

### Accuracy comparison with alternative solutions

To assess whether using SpaRC improves recovery of the known synthetic spike-in transcripts in the PacBio human data, clustering results from SpaRC were compared with the minimap-based [20] clustering results and run through the PacBio Iso-Seq clustering pipeline [27]. The results (Table 3) from SpaRC show comparable results with slightly improved recovery of the synthetic spike-in transcripts (more true positives) and slightly reduced artifacts (fewer false positives). The difference seems to be more pronounced when sequence depth is lower (PacBio2).

**Table 3:** Comparison of recovered synthetic SIRV transcripts in the PacBio
human transcriptome data.

| Dataset | SpaRC |  | Minimap |  |
| --- | --- | --- | --- | --- |
|  | TP | FP | TP | FP |
| PacBio2 (375k reads) | 57 | 13 | 54 | 17 |
| PacBio3 (623k reads) | 61 | 11 | 61 | 14 |

### Data Complexity has a major effect on SpaRC execution time

As introduced above, SpaRC consists of 4 steps: KMR, GCER, Graph Partition, and AddSeq. To measure the computing efficiency of each step on data with different complexity (number of species, see Table 2), we run SpaRC against Human Alzheimer transcriptome, Maize transcriptome, and Cow Rumen metagenome with the same data size (about 4GB) on OTC.

We found different datasets give rise to very different execution times (Fig 2). First of all, complex metagenome dataset has many more unique kmers, which requires longer KMR running time. Same sized transcriptome dataset, Pabio has more k-mers than Illumina, presumably due to higher error rate. Second, reads from complex metagenome dataset typically have fewer edges than transcriptome because many species do not have sufficient sequencing coverage. Longer reads tend to have more edges because they have more k-mers (Table 4). Finally, even given comparable number of total edges (Table 4), LPA step takes significantly longer execution time for long read transcriptome dataset than the short read transcriptome dataset. This is because each long read dataset has more edges per vertex than short read, and GraphX’s LPA implementation uses vertex-cut for graph partition [38], resulting in more copies of vertices, which in turn translates into higher time cost in each LPA iteration.

**Figure 2.**
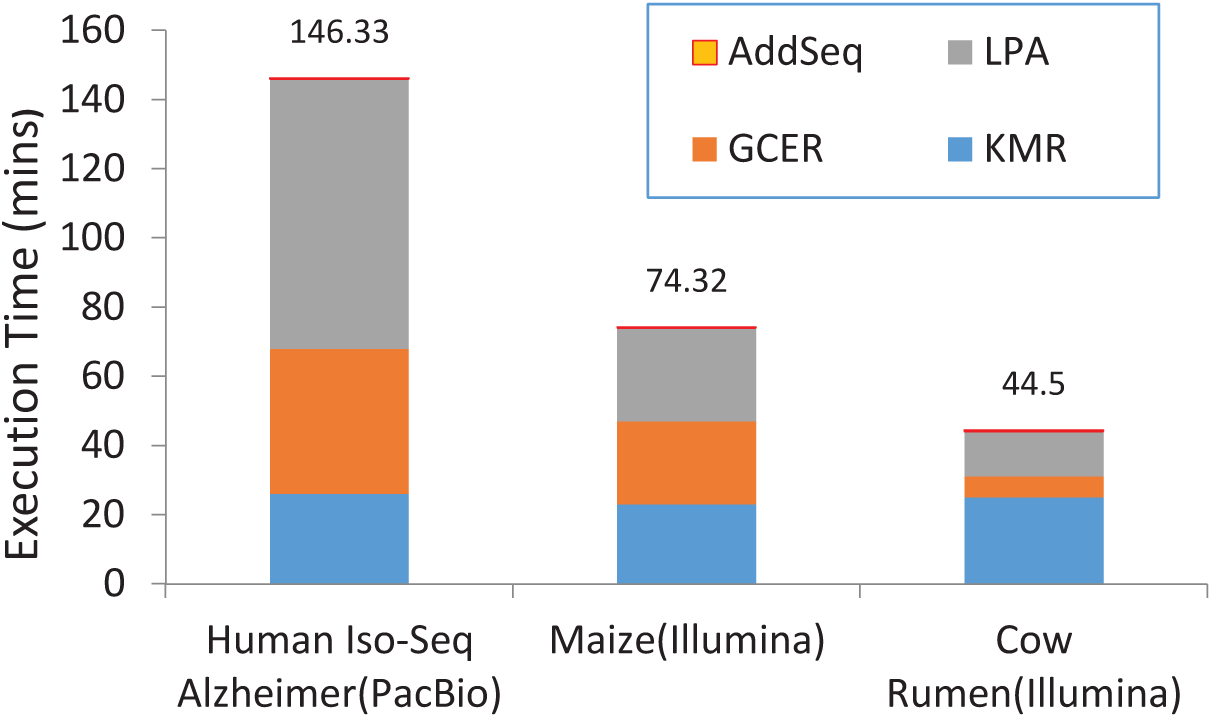
Performance difference between datasets

**Table 4.** Metrics of different datasets

| Dataset | # of k-mers | # of edges | # of nodes | Avg degrees per node |
| --- | --- | --- | --- | --- |
| PacBio1 | 179,039,835 | 263,116,527 | 1,027,204 | 512 |
| Maize | 96,643,966 | 298,631,852 | 11,465,314 | 52 |
| Cow Rumen | 46,027,775 | 41,155,061 | 4,001,389 | 20 |

Among the steps in the workflow, AddSeq is the simplest step and takes very little time (no more than 1 minute) for all datasets.

### Degree of Parallelism

On SpaRC’s computing performance

It has been reported parallelism level has a major effect on the performance of the Spark applications [1]. In earlier versions of SpaRC based on Spark version 1.6, we also observed smaller parallelism level led to poor performance due to some jobs take too long to finish due to data imbalance (data not shown). We therefore evaluated the effect of parallelism level on the overall execution time of the current SpaRC software. Because the size of each data partition is also a function of input data size, we run multiple SpaRC experiments over 20GB and 50GB Cow Rumen dataset on OTC, each with a Spark default parallelism (spark.default.parallelism) value ranging from 50 to 20,000. Once set, Spark automatically sets the number of partitions of an input file according to its size for distributed shuffles.

As shown in Fig 3, we found the performance of SpaRC does not vary much over several orders of magnitude in parallelism, for both of the two datasets tested. As long as the parallelism is not extreme (less than 100 or over 1 million), SpaRC’s performance is quite consistent. When there are too few data partitions, performance suffers because of cluster resource under utilization. In contrast, when there are too many data partitions, there might be excessive overhead in managing small tasks. It is not necessary, at least in this case, to adjust the default parallelisms.

**Figure 3.**
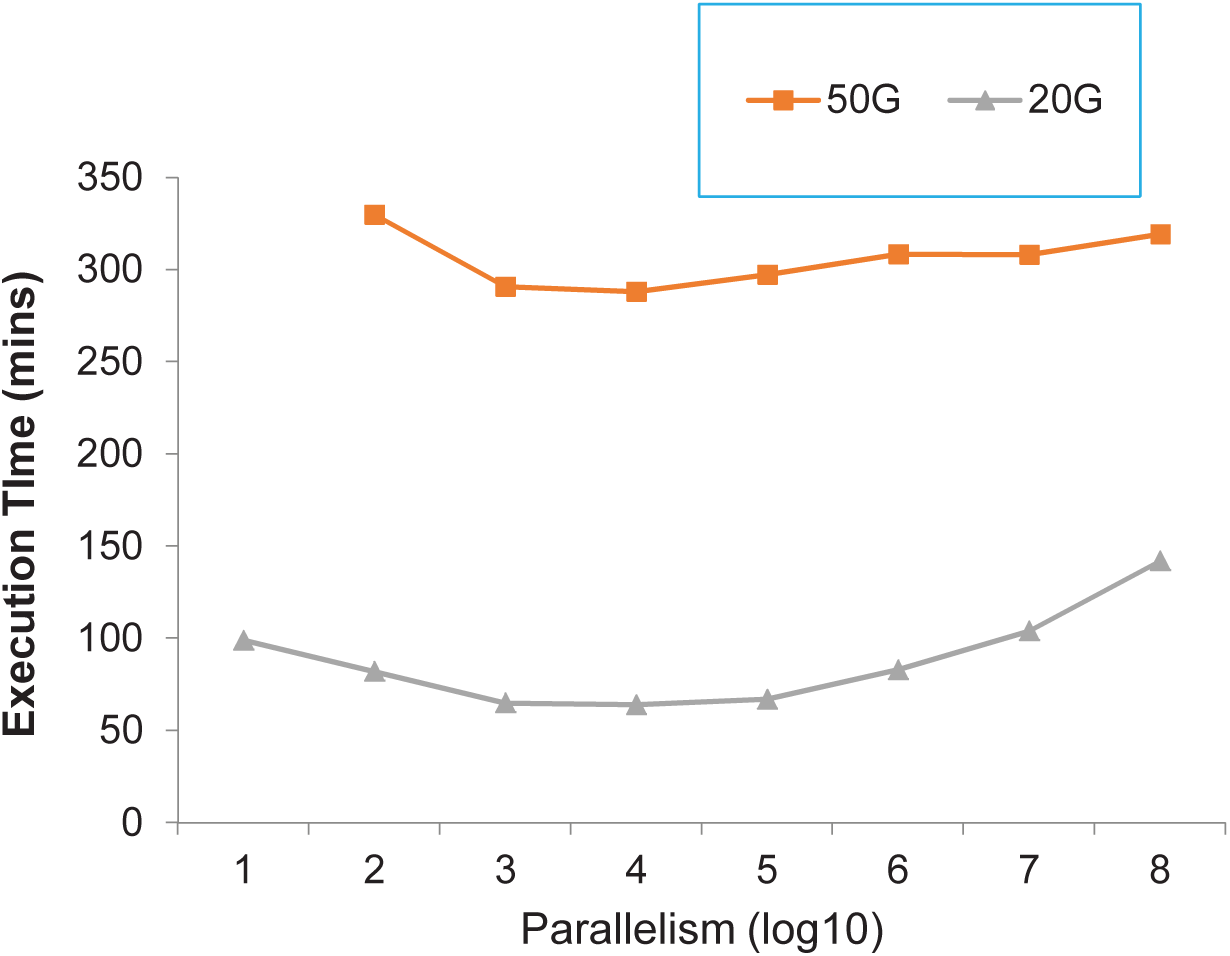
The effect of parallelism level on the total execution time and GCER Step scale up linearly as expected, while LPA step scales up near linearly, consistent with previously reported [30].

It is worth noting that Spark relies on Hadoop file system (HDFS) which has a default partition size 64MB. Our previous work showed that bioinformatics applications can benefit from setting it to 32MB [32], therefore in SpaRC we recommend setting HDFS default partition size to 32MB.

### SpaRC scales near linearly with input data and compute nodes

We designed two different experiments to measure the scalability of the SpaRC. The first one tests its data scalability as more input are added on a fixed-sized cluster, and the second measures its horizontal scalability as more nodes are added to the cluster to compute the same input. For data scalability test we use 20GB, 40GB, 60GB, 80GB, and 100GB fastq datasets from Cow Rumen metagenome. The sequence retrieval step (AddSeq) is not shown due to its negligible processing time (as mentioned in the above).

We report in Fig 4 the result of the first experiment varying input data size and maintaining the number of nodes in the OTC cluster to a fixed value (20). The KMR

**Figure 4.**
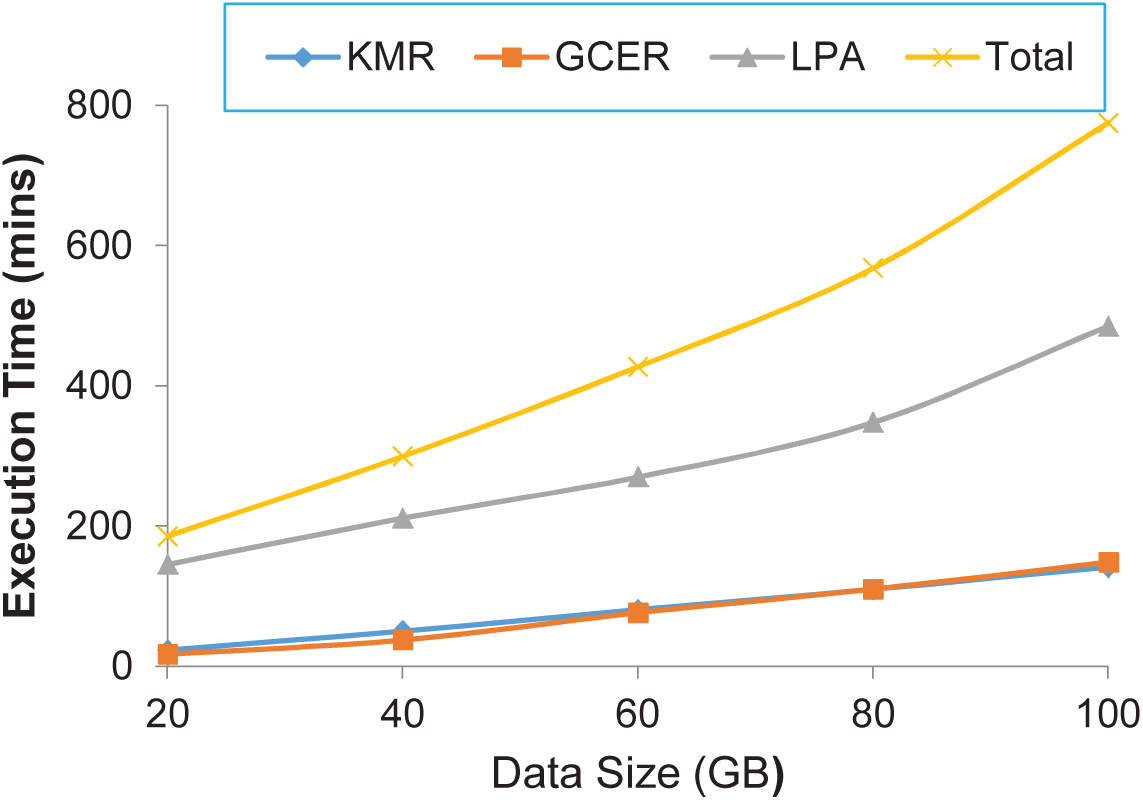
Scalability with different input size

We next tested SpaRC performances by keeping the input size fixed (10GB, 50GB) but varying number of nodes. As shown in Fig 5, the compute time required for each stage and the total time decreases as the number of nodes increases. However, there appears to be a “sweet spot” for each specific input size (10 nodes for 10GB, 50 for 50GB, respectively). Before the number of nodes reaches this spot, every doubling in number of nodes translates into approximately halving the compute time. However, the slope of time saving is decreasing when the node number increases beyond the spot.

**Figure 5.**
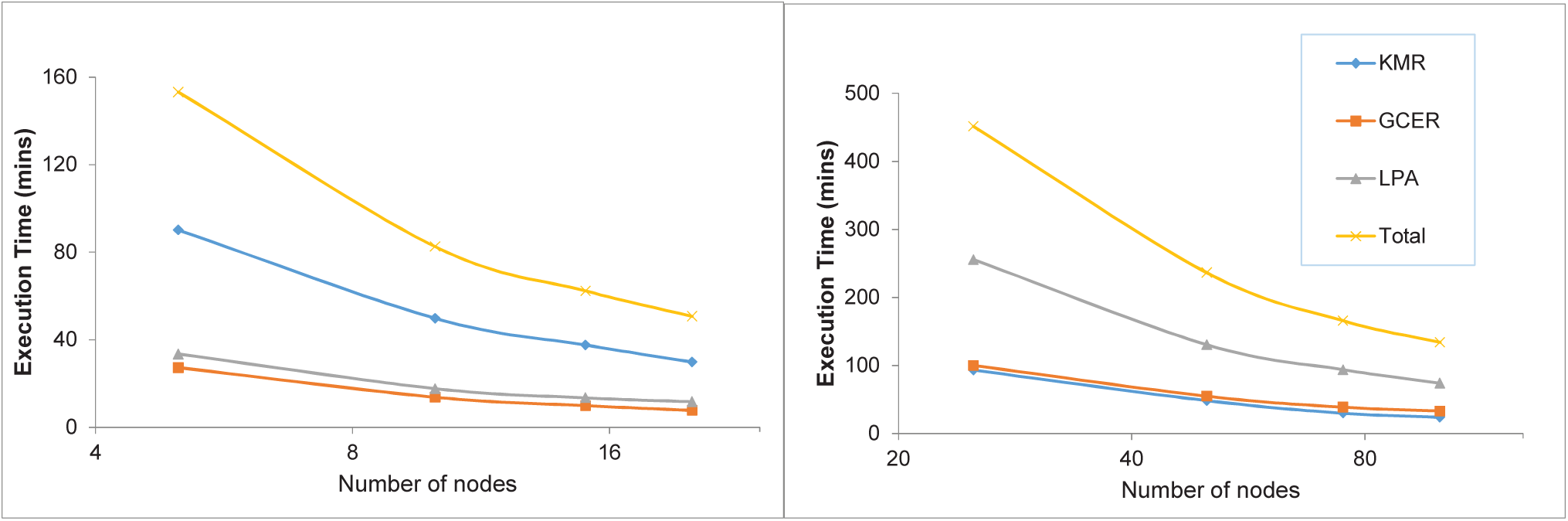
scalability with different number of nodes (Left: 10GB; Right: 50GB)

This phenomenon can be explained by the Amdahl’s law [3] in parallel computing. Overall, we achieve the near-linear scalability as other spark-based tools [8,31], suggesting SpaRC scales well to the number of nodes.

## Conclusion and Discussion

Metagenome and transcriptome assembly is challenging due to both its scale and complexity. Here we developed a scalable algorithm, SpaRC, for large-scale metagenome and transcriptome reads clustering to enable downstream assembly optimization tailored towards individual gene/genome. SpaRC takes advantage of Apache Spark for scalability, efficiency, fast development and flexible running environments. We evaluated SpaRC on both transcriptome and metagenome datasets and demonstrated that SpaRC produces accurate results comparable to state-of-the-art clustering algorithms.

Since Apache Spark is still a very young project undergoing heavy development, some of its components have not been stable and/or optimized. For example, currently LPA is implemented in GraphX using the pregel interface [21] instead of in GraphFrame [10], which did not take the full advantage of the scalability and efficiency of DataFrame API [6]. Current LPA function in GraphFrame is a simple wrapper of the method in GraphX, and it is neither space nor time efficient. Since it cumulatively caches the results of each iteration for job recovery, disk usage often explodes as the number of iterations increases. Furthermore, if one executor dies, all of its cached data is lost and the whole process has to start from scratch. Creating a checkpoint for each iteration like the GraphFrame version of connect component should alleviate this problem.

We observed the clusters produced tend to be too small when the read length is short (e.g., single-end metagenomic dataset on Illumina platform). For pair-end sequencing datasets one can merge (if they overlap) or concatenate the two ends to increase the cluster size. Decreasing k-mer size, or requiring less shared k-mers should also help increase cluster size. However, this may lead to decrease of purity. One potential solution is to run an additional binning or scaffolding step (using pair-end or long reads if available) after assembling each cluster of reads into contigs, a common step in metagenome assemblies.

Based on our experience running SpaRC on OTC and AWS cloud computing environments give similar performance. We also attempted to run SpaRC on HPC environments, including NERSC’s Cori system (http://www.nersc.gov/users/computational-systems/cori/) and Pittsburgh Supercomputing Center’s Bridge system (https://www.psc.edu/bridges). On these systems, the Hadoop and Spark frameworks are provisioned in an on-demand fashion to allow Spark jobs. Although SpaRC runs well on small datasets on both systems, scaling up to larger dataset (>1Gb) failed because of various job scheduling and network problems. More research is needed to get SpaRC run in similar HPC environments.

## Acknowledgments

We thank Drs XXX for critical reading the manuscript. We thank members of NERSC, especially Dr. Lisa Gerhardt and Mr. Evan Racah for their support to run SpaRC on Cori system. We thank members of Pittsburgh Supercomputing Center (PSC), especially Dr. Philip Blood and Mr. Bryon Gill for their support to run SpaRC on the Bridge system. We thank Dr. Hui Zheng and Mr. Billy Xu from Huawei Inc. for their support to run SpaRC on the OTC system.

## Funding

Xiandong Meng and Zhong Wang’s work was supported by the U.S. Department of Energy, Office of Science, Office of Biological and Environmental Research under Contract No. DE-AC02-05CH11231. The OTC computing resource is provided by Huawei Inc. through research collaboration. Part of the Amazon Web Service computational resources was supported by the AWS Cloud Credits for Research Program “EDU_R_RS FY2015 Q4 DOI-JointGenomeInstitute Wang”. Computing resource on PSC’s Bridge system was supported by an XESED grant no “MCB170069”.

